# Machine-learning-based Structural Analysis of Interactions between Antibodies and Antigens

**DOI:** 10.1101/2023.12.06.570397

**Authors:** Grace Zhang, Zhaoqian Su, Tom Zhang, Yinghao Wu

## Abstract

Computational analysis of paratope-epitope interactions between antibodies and their corresponding antigens can facilitate our understanding of the molecular mechanism underlying humoral immunity and boost the design of new therapeutics for many diseases. The recent breakthrough in artificial intelligence has made it possible to predict protein-protein interactions and model their structures. Unfortunately, detecting antigen-binding sites associated with a specific antibody is still a challenging problem. To tackle this challenge, we implemented a deep learning model to characterize interaction patterns between antibodies and their corresponding antigens. With high accuracy, our model can distinguish between antibody-antigen complexes and other types of protein-protein complexes. More intriguingly, we can identify antigens from other common protein binding regions with an accuracy of higher than 70% even if we only have the epitope information. This indicates that antigens have distinct features on their surface that antibodies can recognize. Additionally, our model was unable to predict the partnerships between antibodies and their particular antigens. This result suggests that one antigen may be targeted by more than one antibody and that antibodies may bind to previously unidentified proteins. Taken together, our results support the precision of antibody-antigen interactions while also suggesting positive future progress in the prediction of specific pairing.

## 1. Introduction

Antibodies are the central player in the humoral immune response, in which these proteins are secreted by B lymphocytes to capture the foreign or disease-associated antigens [1]. The recognition of antigens by antibodies is achieved through their highly specific interactions [2]. These interactions are mediated by the residues located on the binding interfaces of antibody-antigen complexes. The residues on the interface of antigens are called epitopes, while the residues on the antibodies, mainly situated in their complementarity-determining regions (CDRs), are known as paratopes [3]. A long-standing but still unsolved question is whether we can predict the pairing between a specific paratope and its corresponding epitope. Studies toward this direction provide the basis for the functional characterization of antibody-antigen interactions, which can greatly facilitate our understanding of the molecular mechanism underlying humoral immunity. Moreover, analysis of paratope-epitope interactions can further boost the design of new monoclonal antibodies (MAbs) [4], which are the highly promising therapeutics for many diseases including melanoma [5] and chronic lymphocytic leukemia (CLL) [6].

Epitopes in a protein can be identified by techniques such as phage display libraries [7] and peptide microarrays [8]. The binding between antibodies and antigens can also be inferred by hydrogen/deuterium exchange (HDX) experiments [9]. Comparing with these approaches, computational modeling is much less time-consuming and labor-intensive, and has already been widely applied to study antibody-antigen interactions. Early computational methods used various propensity scores to search epitopes in protein sequences and structures [10, 11]. With known structures of antibody-antigen complexes, the dynamic properties of the interactions between epitopes and paratopes can be estimated by molecular dynamics simulations [12-16]. If their structures are unknown, the antibody-antigen interactions can be predicted by molecular docking [17-22] or homology modeling [23-26]. Given the accumulating experimental data on antibody-antigen complexes in the protein data bank (PDB) and the recent breakthrough in artificial intelligence, machine-learning or deep-learning-based models have been becoming mainstream in the field [27-31]. For instance, different algorithms, including graph convolutional networks [32], long short-term memory networks [33], and protein language models [30], have been employed for linear and conformational epitope prediction. As another example, AbAdapt integrates AlphaFold structure prediction with rigid docking to generate antibody-antigenspecific features [34]. Nevertheless, detecting antigen-binding sites associated with a specific antibody is currently still a challenging problem under extensive studies.

In this study, we computationally analyzed the structures at the binding interfaces between antibodies and the corresponding antigens by a machine learning model. The analysis is based on a large-scale non-redundant structural database of antibody-antigen complexes. The structural information at the interface of an antibody-antigen complex was encoded into a twodimensional matrix. The matrix was then fed to a convolutional neural network for searching the patterns of antibody-antigen interactions. Based on training and cross-validation, we found that our deep learning model can successfully identify antibody-antigen complexes from general protein-protein complexes with high accuracy. More interestingly, if only using the epitope information, we can still identify antigens from the normal protein binding regions with an accuracy higher than 70%, suggesting that there are unique features on the surface of antigens to be recognized by antibodies. However, our model was not able to predict the pairs between antibodies and their specific antigens, implying that an antigen might be targeted by multiple antibodies, vice versa, antibodies can be repurposed to different proteins which may not be discovered before. In summary, our studies provided a useful tool to characterize the structure of antibody-antigen interactions.

## 2. Methods

### 2.1 Dataset contents

We adopted a large dataset consisting of 1215 pairs of antibody-antigen interactions for computational analysis. The dataset was downloaded from the AbDb database [35]. The threedimensional structure of each complex in the dataset is in the format of Protein Data Bank (PDB). Each complex contains an antibody and its corresponding antigen. The antibody further contains a heavy [H] chain and a light [L] chain, while the antigen could also contain multiple subunits that are marked by other letters. Moreover, AbDb performed pairwise comparisons across the sequences of all antibodies to eliminate redundant information in the database. As a result, all antibody-antigen complexes in our dataset are not identical.

At the same time, a control dataset of general protein-protein interactions was used for comparison. The dataset contains 4960 protein complexes and was constructed by us from the original 3did database [36, 37]. The 3did database includes a large variety of protein complexes for which high-resolution three-dimensional structures are available. We reduced the redundancy in this database by removing the complexes that share high sequence identities with others. Detailed procedure can be found in our previous study [38, 39].

### 2.2 Calculating the feature matrix of protein-protein interactions

The interaction between a pair of antibody and antigen with known three-dimensional structure was converted to a two-dimensional feature matrix as follows. We first extracted all the residues that are located at the binding interface on both antibody and antigen sides. A residue from an antibody is considered to be at the binding interface if it is located in the CDR loops and forms contact with any residue from the antigen. Similarly, a residue from an antigen is considered to be at the binding interface if it forms contact with any residue from the CDR loops of its corresponding antibody. A distance cutoff of 5 Angstrom was used to define the contacts between heavy atoms of the sidechains from two residues [40]. All pairs of intramolecular contacts between interface residues in an antigen were further enumerated using the same distance cutoff. The combinations of residue type for all these intramolecular contacts were transferred into the upper diagonal section of a matrix, as illustrated in **Figure 1**. The dimension of the matrix is 20×20, in which 20 denotes the type of amino acids. The value of a specific matrix unit indicates the frequency of finding the corresponding combination of two residue types at the interface region of the given antigen. Following the same procedure, the residue type combinations of all intramolecular contacts in an antibody were transferred into the lower diagonal section of a matrix. As a result, the full feature matrix of an antibody-antigen complex was constructed by combining the two diagonal sections.

**Figure 1.**
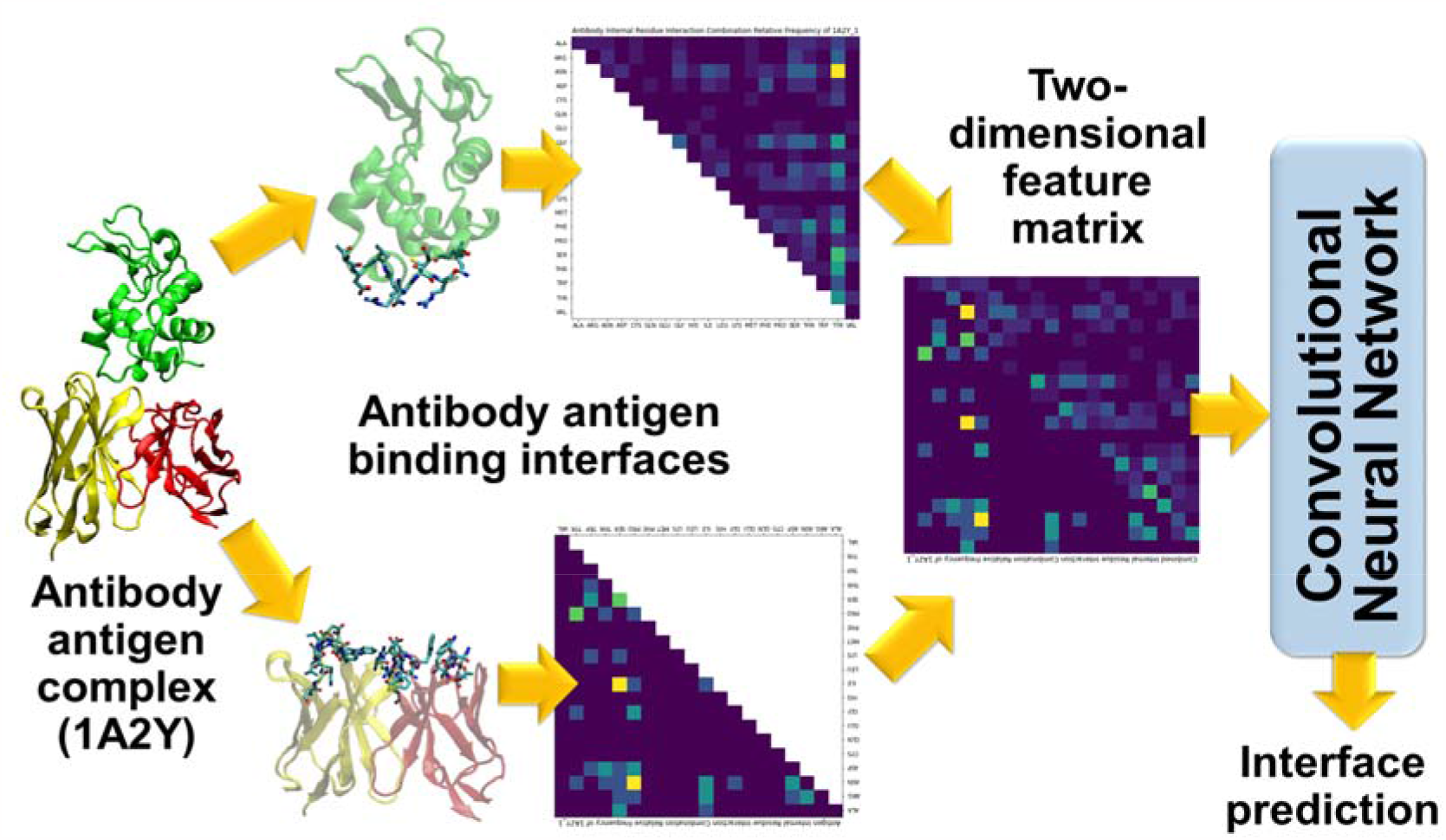
Outline of process. A matrix of antibody and antigen internal interactions with normalized frequency values are created from structural protein data. The combined matrix is used to train the convolutional neural network.

Using this strategy, we calculated the feature matrix for all 1215 complexes in the antibody-antigen dataset, as well as for all 4960 complexes in the general protein-protein interactions dataset.

### 2.3 Machine learning algorithm

As shown in **Figure 1**, the feature matrices of antibody-antigen complexes constructed above were normalized and then fed into a 2D convolutional neural network (CNN). Each of the three layers of the CNN is convolved with 32 filters (3×3 window size). A maximum pooling layer follows each convolutional layer to reduce feature map sizes. The remaining output values from the convolutional layers are passed into a multilayer perceptron (MLP) that has one layer with 512 neurons, a second layer with 64 neurons, and a third layer for output. The MLP reduces the final output into a single category prediction.

Using this CNN classifier, we carried out three different tests. Within each test, a benchmark set was set up by selecting matrices of various antibody-antigen or general proteinprotein interactions as positive and negative data. We applied 10-fold cross validation (scikitlearn library’s StratifiedKFold) to assess the accuracy of each test by shuffling the benchmark set ten times. The CNN is then trained and validated on each set, which is used to generate an average accuracy. More details are described in the results.

### 2.4 Performance calibration

The extent of the CNN’s proficiency after being trained in each test can be determined from the receiver operating characteristic (ROC) curve of the test results. The ROC is generated by plotting the true positive rate (TPR) against the false positive rate (FPR) to show how well the model classifies positive and negative results. First, the model was verified to be able to completely distinguish antibody-antigen pairs from random noise, with an area under the curve (AUC) score of 1. Thus, the ROC AUC is essentially a square and perfectly performs the calibration for distinguishing two classification thresholds. Every test detailed in this study also utilizes two classification groups for the CNN to differentiate between. Once implemented in the actual tests, therefore, the ROC AUCs of the CNN predictions are able to accurately provide a metric to evaluate the model, with a greater ROC AUC score indicating greater accuracy.

## 3. Results

### 3.1 Residues Preference at Interface

Based on the calculations of all antibody-antigen complexes in the dataset, the histogram in **Figure 2** depicts which amino acids are most frequently observed at the binding interfaces.

**Figure 2.**
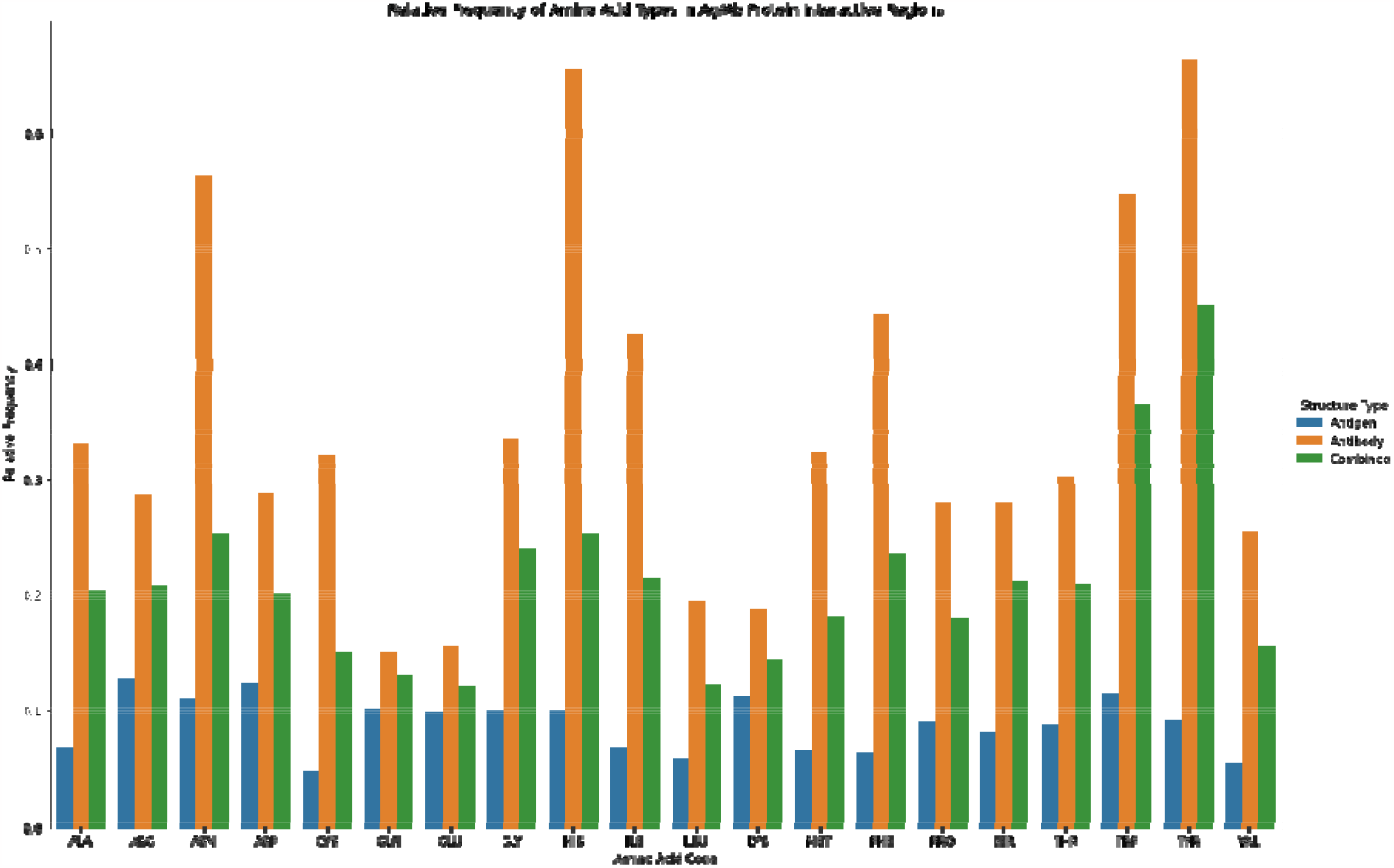
Histogram of amino acid frequencies derived from all 1215 antibodyantigen interactions in the dataset.

This value is determined by having two dimensions, one being a residue from the antibody and the other, a residue from the antigen. The relative frequency is then the prevalence of finding an amino acid pair between the two surfaces.

Generally, the frequency of an amino acid on either the antigen or antibody structure is correlated, as seen in the histogram. For example, if one amino acid’s frequency value on the antibody structure is greater than another amino acid, then its frequency value on the antigen structure will generally also be greater. However, there are discrepancies such as in Tryptophan (TRP) and Tyrosine (TYR) where TRP has a higher frequency value on the antigen structure (0.1149 versus 0.0916) but a lower frequency value on the antibody structure (0.3020 versus 0.5473). The figure shows that TYR and TRP are highly populated in the CDR regions of antiboies, which is consistent with previous observations [41].

The heat map in **Figure 3** contains interactions between the antibody and antigen and the frequency of the amino acid pairs. To create this heat map, all interactions are first counted so the antigen values have their own total for each antibody. Then, each cell is divided along rows and columns twice, once by the antigen total and once by the antibody total. The white value combination (Cysteine, CYS) was originally too high (0.000768) and was consequently decreased to approximately 1.14E-4, originally the second highest value, to make value differences clearer on the gradient scale.

**Figure 3.**
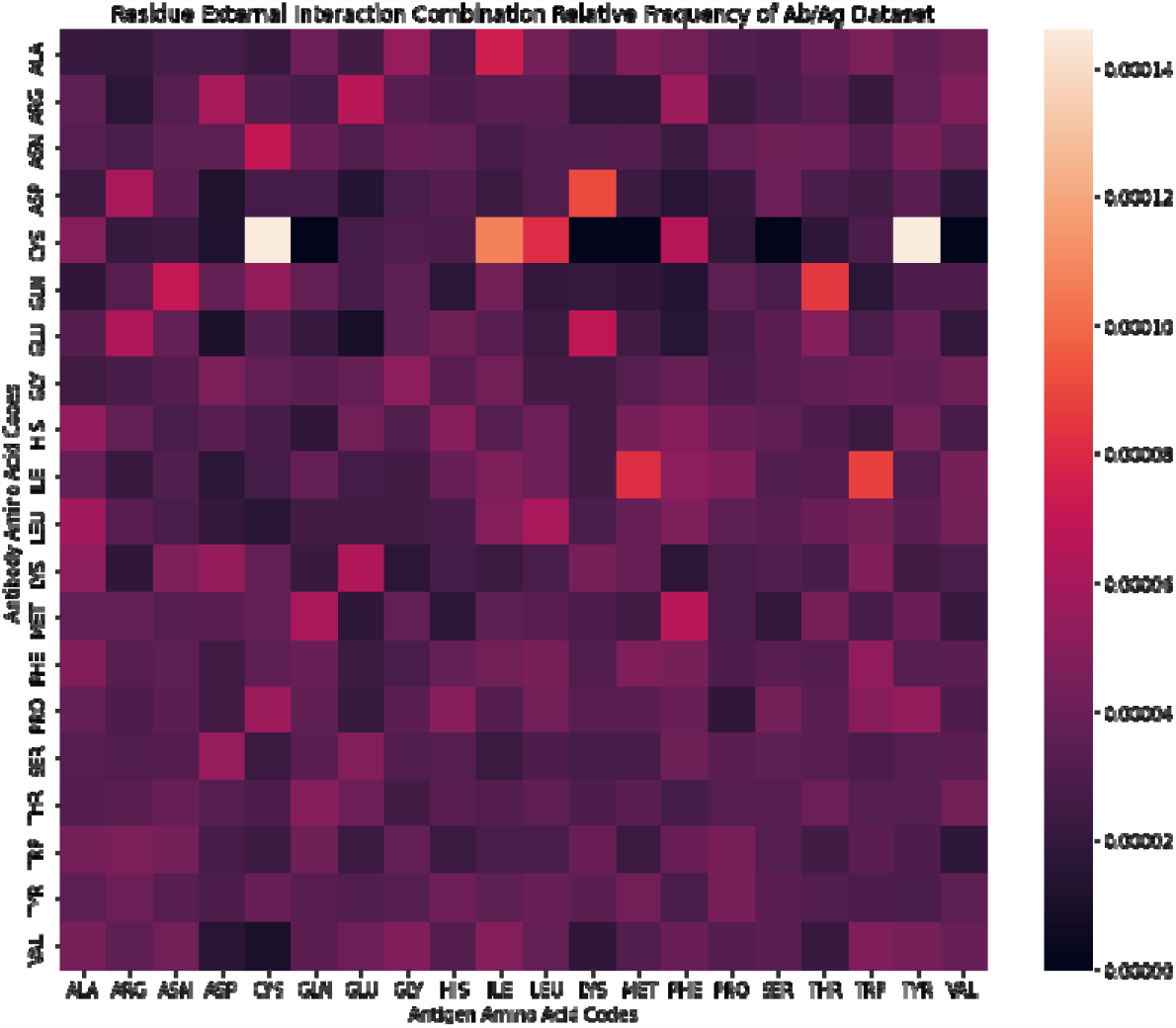
Heat map of antibody-antigen interactions and amino acid pair frequencies.

### 3.2 Distinguish Antibody-Antigen interactions from General Protein-protein interactions

The first testing model used the CNN to compared general protein complexes with antibody-antigen pairs. The goal of this calculation is to have the CNN predict which category an interaction (antibody-antigen interaction or general protein-protein interaction) is when given an input. Feature matrices of internal interaction frequency are created, for the antigen and antibody, respectively. The matrices from each antibody and antigen are combined to represent one complex (**Figure 4**). Points in these charts contain normalized values between 0 (purple and blue) and 1 (yellow), indicating each amino acid pair’s interaction frequency. Some examples of the antibody-antigen interaction matrices are shown in **Figure 5**. Combined feature matrices are then split into training and testing sets for the CNN. Similar matrices are also created for general protein pairs from the second dataset. Similar to the antigen internal interactions, the interactions within these proteins are determined by a set interactive region distance. Some examples of the general protein-protein interaction matrices are shown in **Figure 6**.

**Figure 4.**
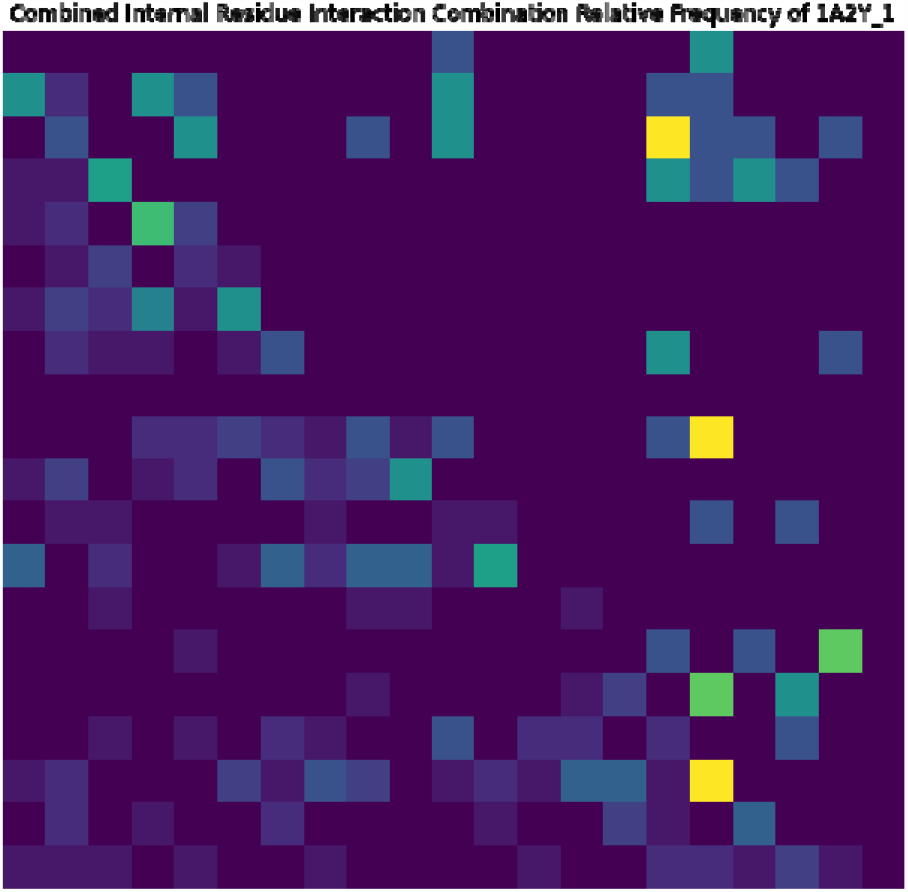
Combined internal interaction frequencies of antibody and antigen into a feature matrix.

**Figure 5.**
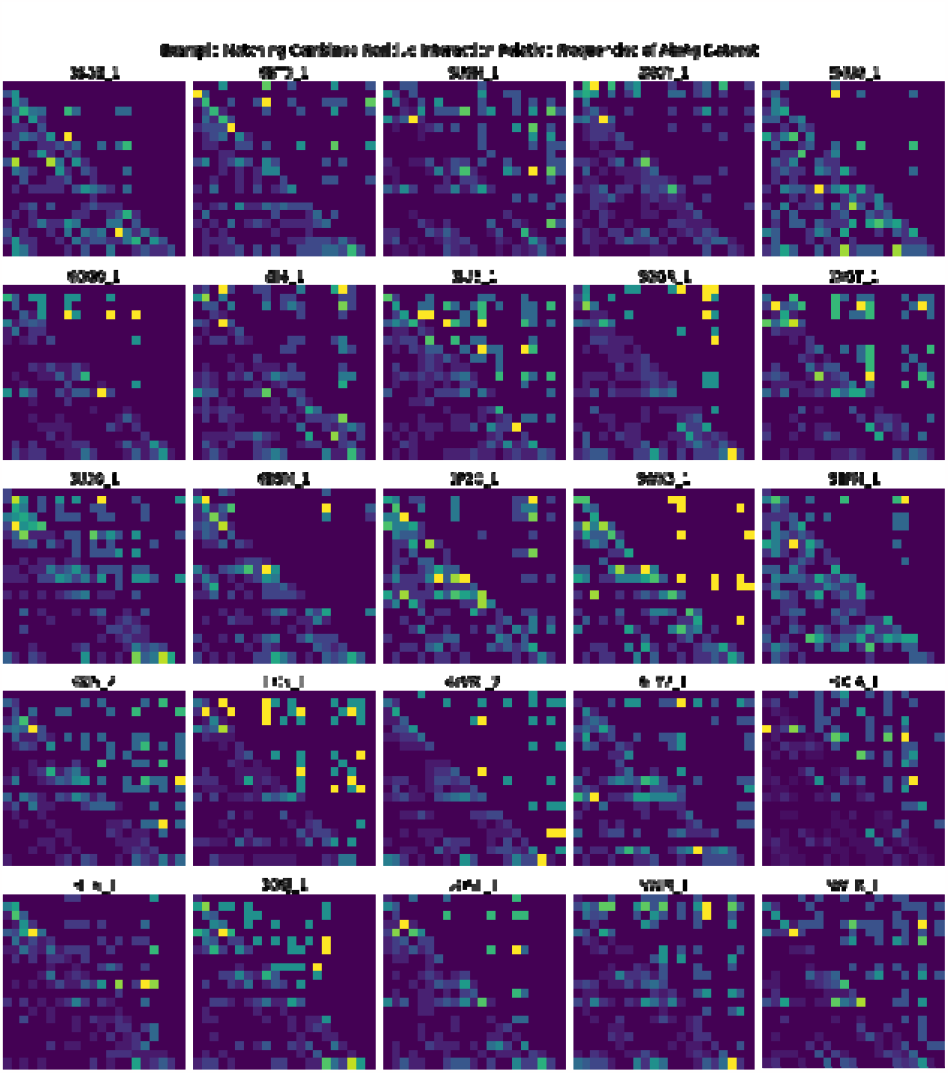
Examples of combined antibody-antigen feature matrices.

**Figure 6.**
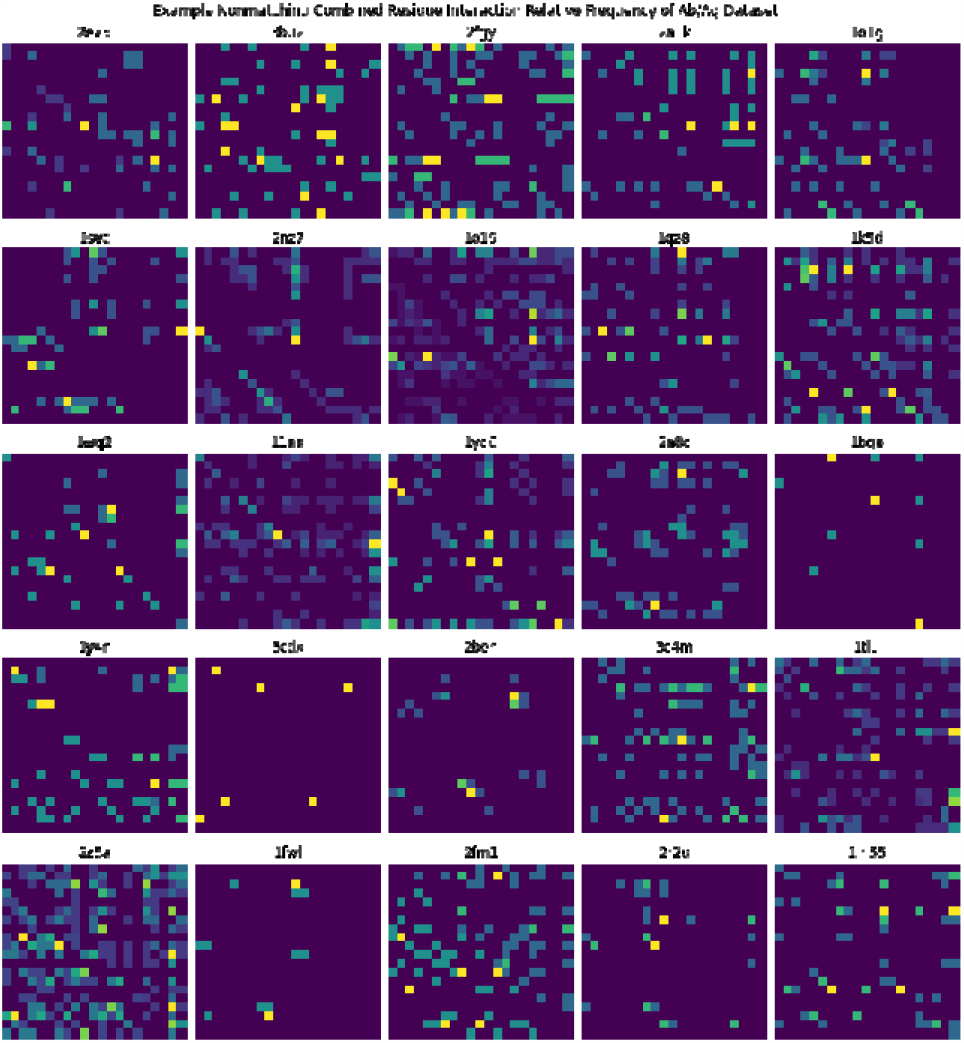
Examples of general protein feature matrices.

In this first test, the CNN’s ability to decide between interactions is checked after being given the training matrixes. Either an antibody-antigen interaction or two interacting chains in general proteins is given to the CNN in testing, and it decides the type of interaction. The average accuracy is determined to be 99.84 percent with the area under the ROC curve (**Figure 7**) being 99.99 percent. The average results from the 10 runs of cross-validation are summarized in **Table 1**. The relatively high accuracy for the CNN on this first test suggests that the antibody-antigen specific interactions can be consistently distinguished from interactions between other types of proteins. The CNN being able to separate the proteins imply a clear difference between the tested matrices of antibody-antigens pairs and general protein pairs, which concurs with the high specificity of protein binding sites. These results indicate positive prospects in the prediction of binding sites due to the ability to separate antibody-antigen pairs by using their interactions.

**Table 1:** The average cross-validation results of antibody-antigen interactions versus general.

| Accuracy | Loss | AUC | Precision | Recall | F1 score |
| --- | --- | --- | --- | --- | --- |
| **0.99842088** | **0.01379772** | **0.99994375** | **0.99672798** | **0.99754099** | **0.99712268** |

**Figure 7.**
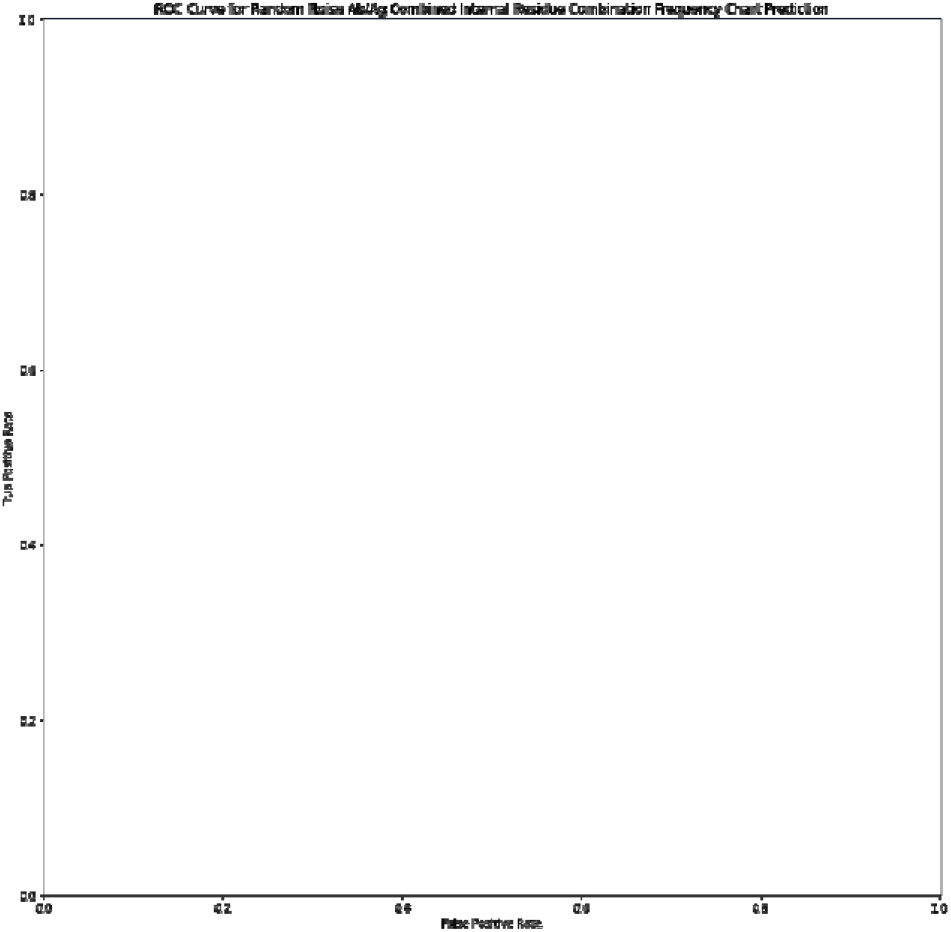
ROC curve for Test 1 (identification of antibody-antigen interactions from general interactions).

### 3.3 Identify epitopes from general protein binding interfaces

The second test assesses accuracy in predicting between general protein interactions and antibody-antigen binding with only the information from antigens. This method utilizes the same distance standards as the first test in order to gauge internal interactions within each antigen. This model trials the extent to which solely antigen internal interactions can estimate types of binding sites and frequency of interactions on the surface of proteins by predicting whether a given protein belong to antigen or other general proteins.

10-fold cross validation of the antigen-only diagonal matrices compared with the general diagonal matrices shows that the average accuracy for this test is 71.11 percent (28.88 percentage points lower than the previous test), with an AUC of 76.00 percent. The ROC curve is plotted in **Figure 8** and the average results from the 10 runs of cross-validation are summarized in **Table 2**. The accuracy of the CNN in differentiating between the antigen-only internal interaction matrices and general protein matrices is lower than in the first test, where the diagonal matrices for both antibodies and antigens were combined for testing. Since only two proteins were individually compared rather than interacting pairs, the CNN prediction results show that internal interactions of antigens versus general proteins differed less from each other. Whereas in the first test the diagonal matrices of antigens and antibodies were combined to show a high specificity in their binding sites, this test only has the individual antibody and general protein. Their internal interactions, therefore, were likely more difficult to separate. However, the accuracy of the CNN’s predictions are still relatively high and suggest that antigen sites can be distinguished.

**Table 2:** The average cross-validation results of antigen binding surfaces versus general binding.

| Accuracy | Loss | AUC | Precision | Recall | F1 score |
| --- | --- | --- | --- | --- | --- |
| **0.71111111** | **0.58215292** | **0.76000542** | **0.69411606** | **0.76529603** | **0.72555238** |

**Figure 8.**
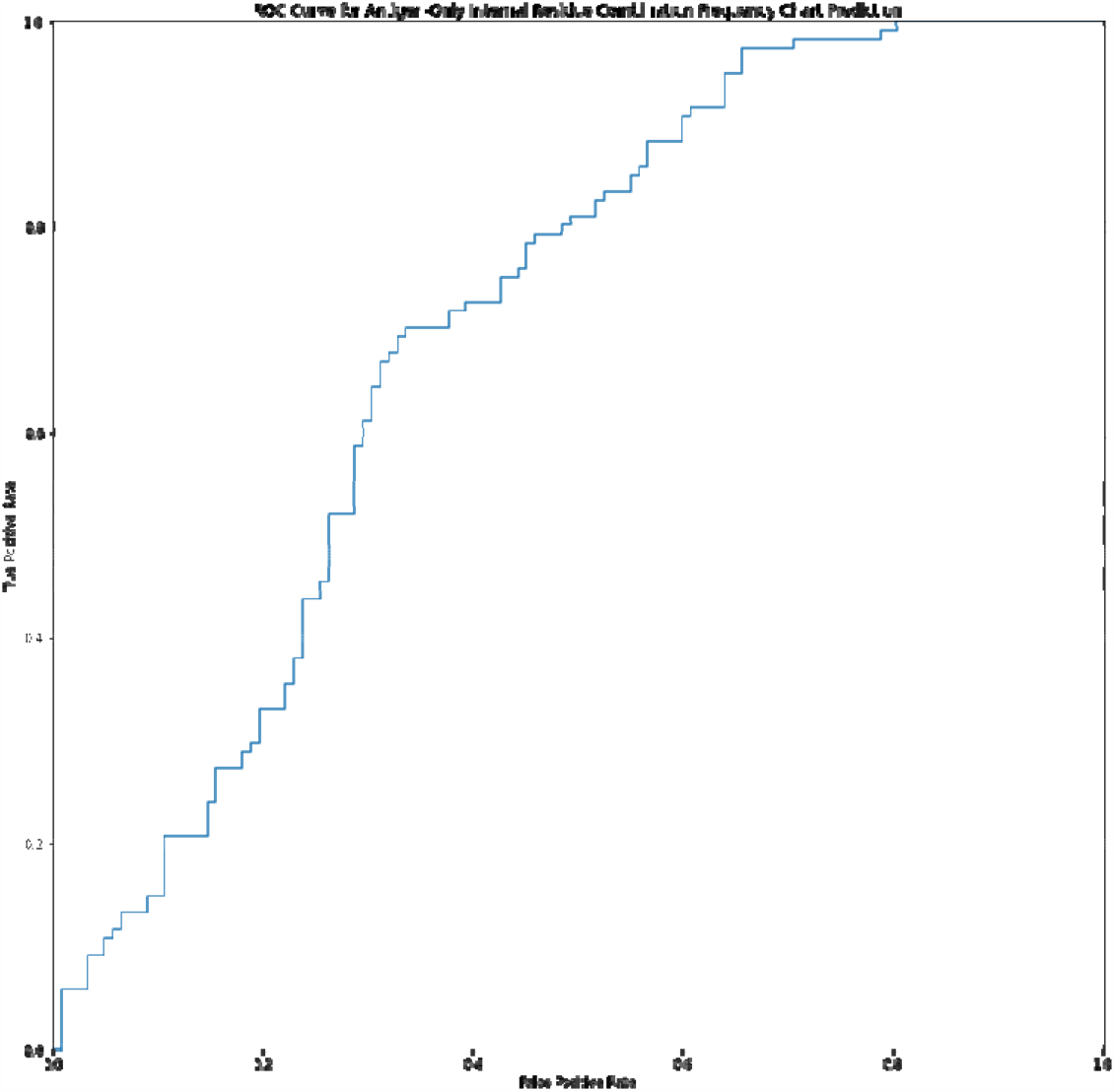
ROC for Test 2 (identification of antigen binding surfaces from general binding surfaces).

### 3.4 Predict interactions between antigens and their specific targeted antibodies

Rather than having two categories of antibody-antigen versus general proteins, this test uses the antibody-antigen group and a second group with mismatched matrices for the machine learning model. The mismatched group has antibodies and antigens that do not coexist in the dataset but are artificially combined together into matrices. The CNN then has to predict whether a given interaction is an actual binding antibody-antigen pair or not. This model has an accuracy of 53.74 percent and AUC of 50.70 percent, lower than both prior tests (testing for antibody-antigen pair versus general protein pair and testing for antigen interaction versus general protein). The ROC curve is plotted in **Figure 9** and the average results from the 10 runs of crossvalidation are summarized in **Table 3**. The lower accuracy of the CNN in this test can be attributed to the greater similarity in training groups. Since both the proper antibody-antigen pairs and the mismatched pairs are of antibodies and antigens, the interaction matrices look more alike to each other than in either previous test. The results of this test convey that the binding sites of each antibody-antigen pair are more difficult to distinguish by the CNN based on interaction matrices despite each protein’s distinct structure.

**Table 3:** The average cross-validation results of real antibody-antigen pairs versus artificially constructed pairs.

| Accuracy | Loss | AUC | Precision | Recall | F1 score |
| --- | --- | --- | --- | --- | --- |
| **0.53744856** | **0.69309179** | **0.50695366** | **0.5386768** | **0.68075464** | **0.57946906** |

**Figure 9.**
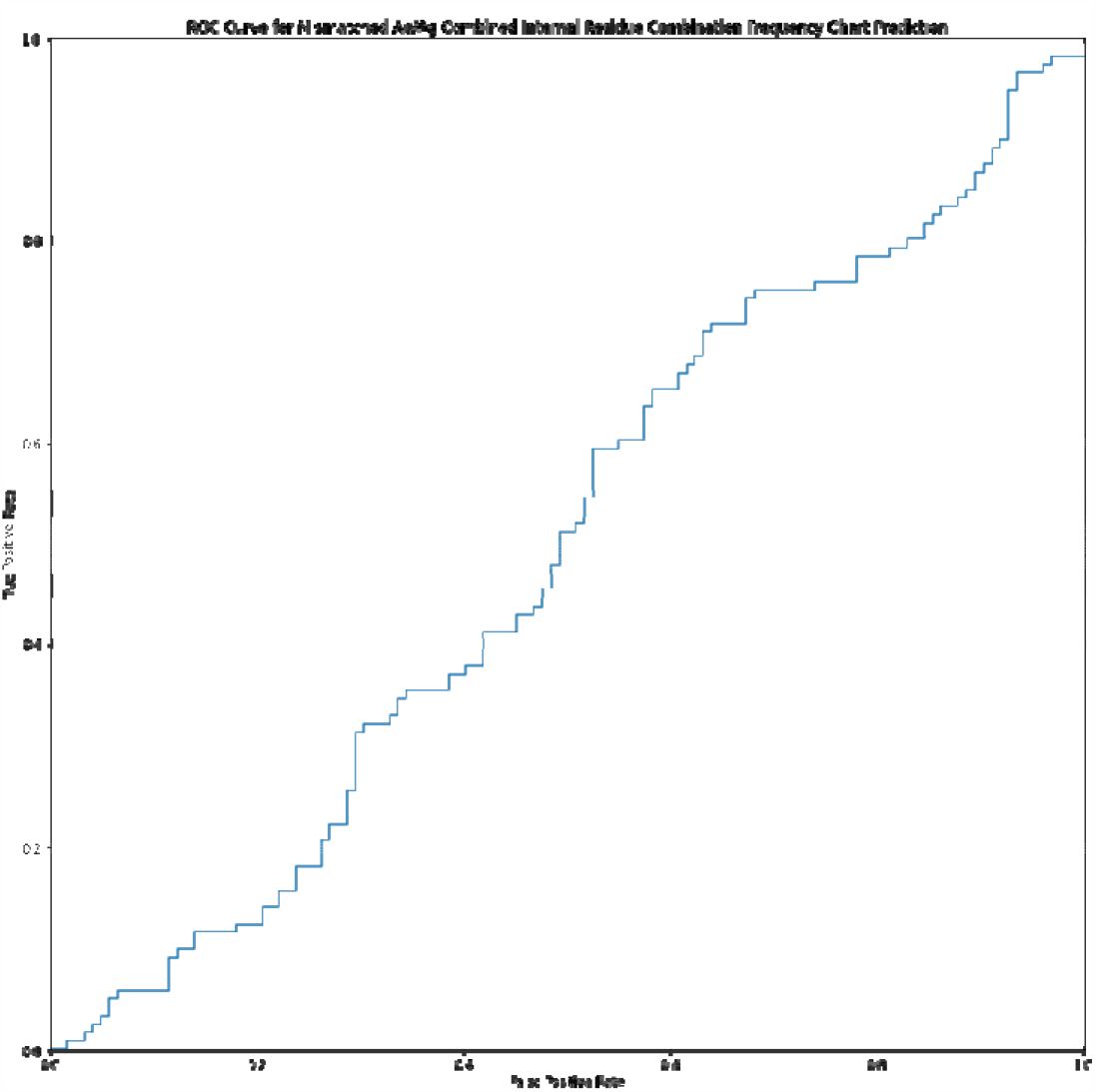
ROC curve for Test 3 (identification of real antibody-antigen pairs from artificially constructed pairs).

## 4. Concluding Discussions

This study trained CNN with groups of antigens, antibodies, and other general proteins, with the final goal being to have the machine learning model distinguish between different types of protein-protein interactions. The test using the features of both antibodies and antigens yielded the most accurate results, while the test using just the antigen binding region frequency yielded less accurate but still positive results. Meanwhile, the dataset of mismatched antibody-antigen pairs led to less clear distinction by the CNN, potentially due to the similarity between antibody-antigen pairs. This result implies that the interaction frequencies within and between antigens and antibodies are not enough for the machine learning model to distinguish the minute nuances of each protein binding. In order to improve the accuracy, the model likely must be given additional features, such as secondary structural types or solvent accessible surface of each residue at the binding interface, in addition to the type of amino acids. However, the fact that the model is unable to identify real antibody-antigen complexes from artificially constructed decoys could also be due to the possibility that in reality, one antigen may be targeted by more than one antibody. On the other hand, antibodies may bind to previously unidentified proteins. As a result, the artificially paired antibodies and antigens in the negative set could form interactions under real circumstances.

Due to the varying nature of each antigen, antibody, and protein’s internal interactions, the calculations taken into consideration were based on interaction ranges of amino acids and general interactive distance, respectively for each protein. These generalizations are a slight limitation of the model due to the efficiency gained. Moreover, taking into account each amino acid interaction within the proteins’ binding regions would likely improve accuracy in the future by accounting for every individual protein and antigen’s internal interactions. Additionally, the different frequencies of amino acids (*Figure 2* and *Figure 3*) in the antibody-antigen dataset may have skewed the training of the CNN, because certain amino acids that occur more often on both structures such as Histidine (HIS) and Tyrosine (TYR) could be weighted more in training data. Scaling the dataset by changing the range before applying the machine learning model, similarly to the method in which the heat maps were created, could lead to less bias in the CNN. By implementing a procedure such as inverse scaling, the weights of each amino acid would be better balanced. Consequently, the CNN would focus more on the patterns in frequency on the training heat maps instead of focusing on the specific amino acid combinations that appear more.

## Acknowledgement

This work was supported by the National Institutes of Health under Grant Numbers R01GM120238 and R01GM122804. The work is also partially supported by a start-up grant from Albert Einstein College of Medicine. Computational support was provided by Albert Einstein College of Medicine High Performance Computing Center.

## Author Contributions

G.Z., Z.S., T.Z. and Y.W. initiated the research; G.Z., Z.S., T.Z. and Y.W. designed the research; G.Z., T.Z. and Y.W. performed the research; G.Z. and T.Z. analyzed the data; G.Z. drafted the paper; G.Z. and Y.W. revised the paper.

## Data Availability

All relevant source codes of the CNN models can be found in the GitHub repository: https://github.com/EngiTom/2023AntibodyDataAnalysis.

## References

1. Cyster, J.G. and C.D.C. Allen, B Cell Responses: Cell Interaction Dynamics and Decisions. Cell, 2019. 177(3): p. 524–540.

2. Sela-Culang, I., V. Kunik, and Y. Ofran, The Structural Basis of Antibody-Antigen Recognition. Frontiers in Immunology, 2013. 4.

3. Ramaraj, T., et al., Antigen-antibody interface properties: composition, residue interactions, and features of 53 non-redundant structures. Biochim Biophys Acta, 2012. 1824(3): p. 520–32.

4. Zeng, X., et al., Recent Progress in Antibody Epitope Prediction. Antibodies (Basel), 2023. 12(3).

5. Bhandaru, M. and A. Rotte, Monoclonal Antibodies for the Treatment of Melanoma: Present and Future Strategies. Methods Mol Biol, 2019. 1904: p. 83–108.

6. Christian, B.A. and T.S. Lin, Antibody therapy for chronic lymphocytic leukemia. Semin Hematol, 2008. 45(2): p. 95–103.

7. Rowley, M.J., K. O’Connor, and L. Wijeyewickrema, Phage display for epitope determination: a paradigm for identifying receptor-ligand interactions. Biotechnol Annu Rev, 2004. 10: p. 151–88.

8. Buus, S., et al., High-resolution mapping of linear antibody epitopes using ultrahigh-density peptide microarrays. Mol Cell Proteomics, 2012. 11(12): p. 1790–800.

9. Chalmers, M.J., et al., Differential hydrogen/deuterium exchange mass spectrometry analysis of protein-ligand interactions. Expert Rev Proteomics, 2011. 8(1): p. 43–59.

10. Saha, S. and G.P.S. Raghava. BcePred: Prediction of Continuous B-Cell Epitopes in Antigenic Sequences Using Physico-chemical Properties. in Artificial Immune Systems. 2004. Berlin, Heidelberg: Springer Berlin Heidelberg.

11. Manavalan, B., et al., iBCE-EL: A New Ensemble Learning Framework for Improved Linear B-Cell Epitope Prediction. Front Immunol, 2018. 9: p. 1695.

12. Frost, C.V. and M. Zacharias, From monomer to fibril: Abeta-amyloid binding to Aducanumab antibody studied by molecular dynamics simulation. Proteins, 2020. 88(12): p. 1592–1606.

13. Frota, N.F., et al., Alemtuzumab scFv fragments and CD52 interaction study through molecular dynamics simulation and binding free energy. J Mol Graph Model, 2021. 107: p. 107949.

14. Koçer, İ. and E. Çelik, In silico analysis of the different variable domain oriented singlechain variable fragment antibody-antigen complexes. J Biomol Struct Dyn, 2023: p. 1–11.

15. Margreitter, C., et al., Antibody humanization by molecular dynamics simulations-in-silico guided selection of critical backmutations. J Mol Recognit, 2016. 29(6): p. 266–75.

16. Zhao, J., R. Nussinov, and B. Ma, The Allosteric Effect in Antibody-Antigen Recognition. Methods Mol Biol, 2021. 2253: p. 175–183.

17. Ambrosetti, F., et al., Modeling Antibody-Antigen Complexes by Information-Driven Docking. Structure, 2020. 28(1): p. 119–129.e2.

18. Guest, J.D., et al., An expanded benchmark for antibody-antigen docking and affinity prediction reveals insights into antibody recognition determinants. Structure, 2021. 29(6): p. 606–621.e5.

19. Kilambi, K.P. and J.J. Gray, Structure-based cross-docking analysis of antibody-antigen interactions. Sci Rep, 2017. 7(1): p. 8145.

20. Myung, Y., D.E.V. Pires, and D.B. Ascher, CSM-AB: graph-based antibody-antigen binding affinity prediction and docking scoring function. Bioinformatics, 2022. 38(4): p. 1141–1143.

21. Pedotti, M., et al., Computational docking of antibody-antigen complexes, opportunities and pitfalls illustrated by influenza hemagglutinin. Int J Mol Sci, 2011. 12(1): p. 226–51.

22. Weitzner, B.D., et al., Modeling and docking of antibody structures with Rosetta. Nat Protoc, 2017. 12(2): p. 401–416.

23. Bansia, H. and S. Ramakumar, Homology Modeling of Antibody Variable Regions: Methods and Applications. Methods Mol Biol, 2023. 2627: p. 301–319.

24. Desta, I.T., et al., Mapping of antibody epitopes based on docking and homology modeling. Proteins, 2023. 91(2): p. 171–182.

25. Kuroda, D., et al., Computer-aided antibody design. Protein Eng Des Sel, 2012. 25(10): p. 507–21.

26. Sivasubramanian, A., et al., Toward high-resolution homology modeling of antibody Fv regions and application to antibody-antigen docking. Proteins, 2009. 74(2): p. 497–514.

27. Akbar, R., et al., In silico proof of principle of machine learning-based antibody design at unconstrained scale. MAbs, 2022. 14(1): p. 2031482.

28. Davila, A., et al., AbAdapt: an adaptive approach to predicting antibody-antigen complex structures from sequence. Bioinform Adv, 2022. 2(1): p. vbac015.

29. Huang, Y., Z. Zhang, and Y. Zhou, AbAgIntPre: A deep learning method for predicting antibody-antigen interactions based on sequence information. Front Immunol, 2022. 13: p. 1053617.

30. Shashkova, T.I., et al., SEMA: Antigen B-cell conformational epitope prediction using deep transfer learning. Front Immunol, 2022. 13: p. 960985.

31. Yang, Y.X., P. Wang, and B.T. Zhu, Binding affinity prediction for antibody-protein antigen complexes: A machine learning analysis based on interface and surface areas. J Mol Graph Model, 2023. 118: p. 108364.

32. Lu, S., et al., A Structure-Based B-cell Epitope Prediction Model Through Combing Local and Global Features. Front Immunol, 2022. 13: p. 890943.

33. Saka, K., et al., Antibody design using LSTM based deep generative model from phage display library for affinity maturation. Scientific Reports, 2021. 11(1): p. 5852.

34. Xu, Z., et al., Improved Antibody-Specific Epitope Prediction Using AlphaFold and AbAdapt. Chembiochem, 2022. 23(18): p. e202200303.

35. Ferdous, S. and A.C.R. Martin, AbDb: antibody structure database-a database of PDB-derived antibody structures. Database (Oxford), 2018. 2018.

36. Stein, A., A. Céol, and P. Aloy, 3did: identification and classification of domain-based interactions of known three-dimensional structure. Nucleic Acids Res, 2011. 39(Database issue): p. D718–23.

37. Stein, A., A. Panjkovich, and P. Aloy, 3did Update: domain-domain and peptide-mediated interactions of known 3D structure. Nucleic Acids Res, 2009. 37(Database issue): p. D300–4.

38. Dhusia, K., Z. Su, and Y. Wu, Using Coarse-Grained Simulations to Characterize the Mechanisms of Protein-Protein Association. Biomolecules, 2020. 10(7).

39. Su, Z. and Y. Wu, Computational studies of protein-protein dissociation by statistical potential and coarse-grained simulations: a case study on interactions between colicin E9 endonuclease and immunity proteins. Phys Chem Chem Phys, 2019. 21(5): p. 2463–2471.

40. Zhang, Q.C., et al., PredUs: a web server for predicting protein interfaces using structural neighbors. Nucleic Acids Research, 2011. 39(suppl_2): p. W283–W287.

41. Peng, H.P., et al., Antibody CDR amino acids underlying the functionality of antibody repertoires in recognizing diverse protein antigens. Sci Rep, 2022. 12(1): p. 12555.

